# Variance in Variants: Propagating Genome Sequence Uncertainty into Phylogenetic Lineage Assignment

**DOI:** 10.1101/2021.11.30.470642

**Authors:** David Champredon, Devan Becker, Connor Chato, Gopi Gugan, Art Poon

**Author notes:** contributed equally (David Champredon, Devan Becker).

## Abstract

Genetic sequencing is subject to many different types of errors, but most analyses treat the resultant sequences as if they are known without error. Next generation sequencing methods rely on significantly larger numbers of reads than previous sequencing methods in exchange for a loss of accuracy in each individual read. Still, the coverage of such machines is imperfect and leaves uncertainty in many of the base calls. On top of this machine-level uncertainty, there is uncertainty induced by human error, such as errors in data entry or incorrect parameter settings. In this work, we demonstrate that the uncertainty in sequencing techniques will affect downstream analysis and propose a straightforward method to propagate the uncertainty.

Our method uses a probabilistic matrix representation of individual sequences which incorporates base quality scores as a measure of uncertainty that naturally lead to resampling and replication as a framework for uncertainty propagation. With the matrix representation, resampling possible base calls according to quality scores provides a bootstrap- or prior distribution-like first step towards genetic analysis. Analyses based on these re-sampled sequences will include a more complete evaluation of the error involved in such analyses.

We demonstrate our resampling method on SARS-CoV-2 data. The resampling procedures adds a linear computational cost to the analyses, but the large impact on the variance in downstream estimates makes it clear that ignoring this uncertainty may lead to overly confident conclusions. We show that SARS-CoV-2 lineage designations via Pangolin are much less certain than the bootstrap support reported by Pangolin would imply and the clock rate estimates for SARS-CoV-2 are much more variable than reported.

## 1 Introduction

Generating a genetic sequence from a biological sample is a complex process. Nucleic acids must be extracted from the sample while avoiding contamination by foreign material. If working with RNA, then we must use a reverse transcriptase reaction (which has a high base misincorporation rate) to convert the RNA into DNA.

Polymerase chain reaction (PCR) amplification is often employed to enrich the sample for the target of interest. For next-generation sequencing (NGS) protocols, we have to generate a sequencing library for instance by random shearing of nucleic acids into fragments that are ligated onto special “adaptors”. NGS procedures such as sequencing by synthesis suffer from greater error rate relative to conventional Sanger dye-terminator sequencing, although these rates have continued to improve with new technologies (Fuller et al., 2009; Goodwin et al., 2016; Salk et al., 2018). In addition, the short reads produced by NGS platforms need to be aligned — either by alignment against a reference genome, *de novo* assembly, or a combination of the two — to reconstruct a consensus sequence using one or more bioinformatic programs. Errors can be introduced in any one of these steps (Beerenwinkel and Zagordi, 2011; O’Rawe et al., 2015).

In some cases, naturally occurring variation, *i.e*., genetic polymorphisms, or variation induced by experimental error is directly quantified and encoded into the output. For example, mixed peaks in sequence chromatograms produced from dye terminator sequencing by capillary electrophoresis are assigned standard IUPAC codes (*e.g*., Y for C or T) when the base calling program cannot determine which base is dominant (NC-IUB, 1986). Ewing and Green (1998) and Richterich (1998) both argued that estimates of the base call quality, typically quantified as Phred quality scores (*Q* = − 10log_10_ *P*, where *P* is the estimated error probability), can be an accurate estimate of the number of errors that the machines at the time would make. Due to advances in the technology as well as our understanding of the sources and patterns in the sequencing errors, new error quantification methods and adjustments of existing error probabilities have been proposed (Li et al., 2004, 2009b; DePristo et al., 2011). Nevertheless, Phred scores remain the standard means of reporting the estimated error probabilities for current sequencing platforms. Generally, these scores are used to censor the base calls (*i.e*., label them “N” rather than A, T, C or G) if the estimated probability of error exceeds a predefined threshold. It is also common practice to remove the sequence from further analysis if the total number of censored bases exceeds a maximum tolerance; *e.g*., Doronina (2005); Robasky et al. (2014); O’Rawe et al. (2015). Some authors/tools use more sophisticated models, such as Wu et al. (2017) who use statistical models that incorporate read depth to determine a probability of a sequencing error, but still use the resultant reads to form a consensus sequence with no measure of uncertainty. Furthermore, some studies have extended the concept of per-base error probabilities to calculate the joint likelihoods of partial or full sequences. For example, DePristo et al. (2011) and Gompert and Buerkle (2011) incorporate adjusted Phred scores into a likelihood framework to generate more accurate estimates of genetic diversity within a population; this approach has subsequently been used to develop new estimators of genetic diversity (Fumagalli et al., 2013). Kuo et al. (2018) recently used a similar approach to develop a statistical test of whether a given genome sequence is consistent with a specified alternative sequence. In general, the reported error probabilities from NGS technologies are primarily used for filtering low quality sequences and improving alignment algorithms (which both result in a consensus sequence that is assumed to be error-free) or for hypothesis tests concerning small collections (usually pairs) of sequences.

The uncertainty present in the sequences are seldom propagated to downstream analyses. For example, methods for sequence alignment and homology searches generally employ heuristic algorithms that utilize similarity scores that do not explicitly incorporate the probabilities of sequencing errors. Moreover, methods to reconstruct the evolutionary relationships among sequences as a phylogenetic tree tend to interpret ambiguous base calls as completely missing data, although some exceptions are found in the literature, *e.g*., DePristo et al. (2011). This problem is exacerbated when each sequence represents the consensus of diverse copies of a genome, such as rapidly evolving virus populations where genuine polymorphisms are confounded with sequencing error. See Schneider (2002) for more criticisms of the use of consensus sequences, along with visualizations (Schneider and Stephens, 1990, called *sequence logos*) to display the deviations from a consensus.

Though rare, some studies have proposed methods for propagation of uncertainty from one step to later steps of an analysis. O’Rawe et al. (2015) suggest methods for propagation of sequence-level uncertainty into determining whether two subjects have the same alleles, as well as estimating confidence intervals for allele frequencies. Another exception can be found in Kuhner and McGill (2014), who incorporate an assumed or estimated error rate for the entire sequence into the calculation of a phylogenetic tree and found that incorporation of errors makes the inferred branch lengths much closer to the true (simulated) branch lengths. Though they did not use nucleotide-level uncertainty, Gompert and Buerkle (2011) incorporate the coverage of NGS technologies as part of the uncertainty of estimates for the frequency of alleles in a population. Clement et al. (2010) present an alignment algorithm (called GNUMAP) that takes nucleotide-level uncertainty into account. Their method incorporates Position Weight Matrices into a method of scoring multiple possible matches against a reference genome in order to choose the best alignment. These studies are the exceptions, rather than the rules, and their methods have not yet attained widespread use.

We present a simple general-purpose framework that can be incorporated into any analysis of genetic sequence data. This framework involves converting the uncertainty scores into a matrix of probabilities, and repeatedly sampling from this matrix and using the resultant samples in downstream analysis. Unlike likelihood-based approaches, we do not make assumptions about the underlying patterns or distributions in the data. In so doing, we can gain more accurate estimation of the errors at the expense of computation time. Our technique is amenable to quality score adjustments prior to applying our methods. We demonstrate the impact of propagating sequence uncertainty by applying our methods to the problem of classifying SARS-CoV-2 genomes into predefined clusters known as “lineages” (Rambaut et al., 2020), several of which correspond to variants carrying mutations that are known to confer an advantage to virus transmission or infectivity. We also analyse a collection of SARS-CoV-2 sequences to demonstrate that the estimated rate of new mutations is much more variable than studies relying on deterministic sequences would conclude.

## 2 Methods

### 2.1 Probabilistic representation of sequences

Here, we describe two theoretical frameworks to model sequence uncertainty at the *nucleotide level* or at the *sequence level*. In both frameworks, the sequence of nucleotides from a biological sample is not treated as a single unambiguous observation (known without error), but rather as a collection of possible sequences weighted by their probability.

#### 2.1.1 Nucleotide-level uncertainty

To represent the uncertainty at each position along the genome we introduce the following matrix, which we will refer to as a probabilistic sequence and denote 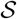:

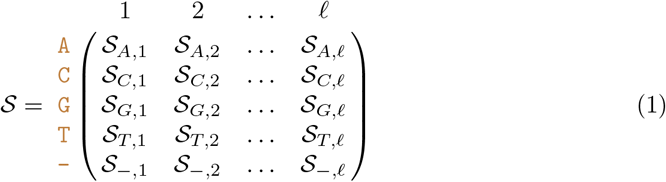

Each column represents a position in a nucleotide sequence of length ℓ. Each row represents one of the four nucleotides `A,C,G,T`, as well as an empty position “−” that symbolizes a recorded deletion rather than missing data. Hence, 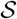 is a 5 × ℓ matrix.

The elements of the probability sequence represent the probability that a nucleotide exists at a given position:

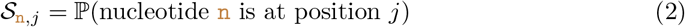

with the special case for a deletion:

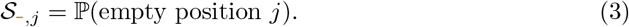

Note that we have for all 1 ≤ *j* ≤ ℓ:

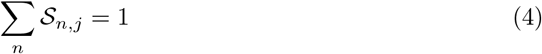

Also, the sequence length is stochastic if 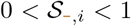 for at least one *i*. The probability that the sequence has the maximum length ℓ is 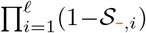. The nucleotide (or deletion) drawn at each position is independent from all the others, so there are up to 5^*ℓ*^ possible different sequences for a given probabilistic nucleotide sequence, but these sequences are *not* equally probable.

A major limitation of this probabilistic representation of a sequence is that we lose all information on linkage disequilibrium. This is especially problematic for recording insertions because insertions with *L* ≥ 2 nucleotides are treated as *L* independent single nucleotide insertions. Instead, we assume that every nucleotide is an independent observation. For example, a probability sequence populated from short read data from a diverse population would not store the information that two polymorphisms were always observed in the same reads, *i.e*., in complete linkage disequilibrium. We also lose information about autocorrelation in sequencing error, such as clusters of miscalled bases associated with later cycles of sequencing-by-synthesis platforms. Sequence chromatograms and base quality scores are affected by the same loss of information.

We note that this representation is similar to the “CATG” file type as described in Kozlov (2018), which indicates the likelihoods of each nucleotide in an aligned mapping for multiple taxa. This file type is able to be used by RAxML-NG to estimate an overall error rate which is then used to estimate phylogenetic trees. Our probability sequence is also similar in concept to Position Weight Matrices (PWMs, Stormo et al., 1982) which are built according to the frequency of each base at each position of a multiple alignment. Our construction differs in that we are creating one matrix per sequence where the entries are weighted according to error probability within that sequence, rather than one matrix for a collection of sequences. However, methods that accept PWMs will be applicable to our probability sequences (and *vice-versa*).

#### 2.1.2 Sequence-level uncertainty

A significant problem of storing probabilities at the level of individual nucleotides is that generating a sequence from this matrix requires drawing ℓ independent outcomes. For example, the reference SARS-CoV-2 genome is 29,903 nucleotides, and a substantial number of naturally-occurring sequence insertions have been described. Thus it would not be surprising if ℓ exceeded 30,000 nucleotides (nt). The majority of these technically possible 5^ℓ^ sequences are not biologically plausible. Therefore, we formulate an ordered subset 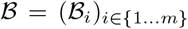 of the first *m* most likely sequences, which are ranked in descending order by the joint probability of nucleotide composition. Note that the sequences in 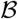, 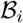, do not necessarily have the same length. The observed genetic sequence, *s*^*^, is a sample from a specified discrete probability distribution *a*:

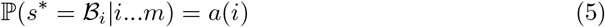

This compact and approximate representation drastically reduces the number of operations to one sample, after some pre-processing to calculate *a*. The observed plurality sequence *s*^*^ (the sequence consisting of the most likely base at each position) is guaranteed to be a member of 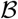 if 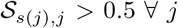 where *s*(*j*) is the *j*-th nucleotide of *s*^*^; indeed, it is guaranteed to be the highest ranked member *i* = 0. We refer to any member of the set 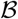 as a *sequence-level probabilistic sequence*. Note that because *a* is a probability distribution, we must have 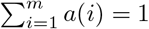. In other words, this probability is conditional on the sequence being in 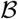.

For example, suppose that we have the following nucleotide-level probabilistic sequence:

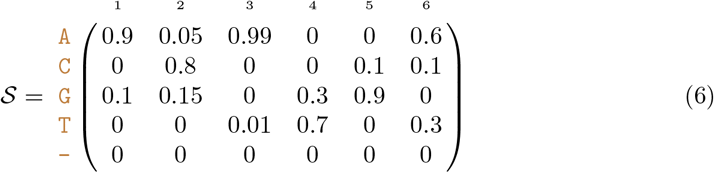

such that there are 2 × 3 × 2^3^ × 3 = 144 possible sequences. The most likely sequence has the highest joint nucleotide probability: `ACATGA` with probability 0.2694 (0.9 × 0.8 × 0.99 × 0.7 × 0.9 × 0.6). If there is a positive probability of deletion for at least one position, then the sequence has a variable length. Large genomes or sequencing targets will result in vanishingly small probabilities for all sequences, and thus calculations on the log scale may be necessary to reduce the chance of numerical underflow.

Table 1 demonstrates the calculation of sequence-level uncertainties using the values in (6). The probability column is the product of the matrix entries for each nucleotide. If the four sequences shown are the only biologically plausible sequences, then the normalized probabilities can be expressed as *α*(*i*).

**Table 1:**
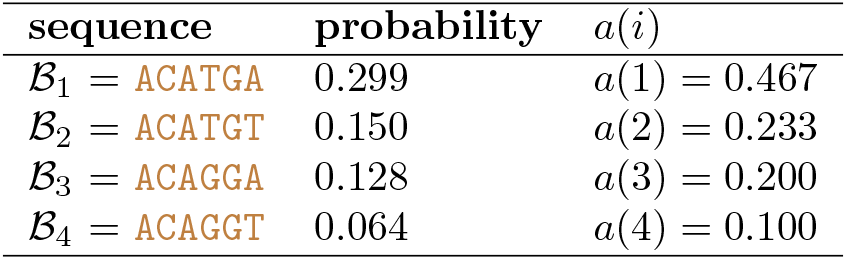
Biologically plausible sequences with probabilities defined by Equation (6)

#### 2.1.3 Deletions and insertions

By construction, the nucleotide-level probabilistic sequence must be defined with its longest possible length. Deletions are naturally modelled with our representation but insertions have to be modelled using deletion probabilities.

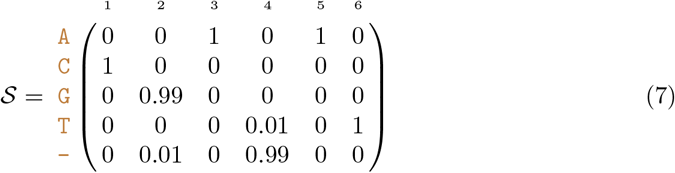

The low deletion probability for position 2 is straightforward to interpret: about 1% of the time, nucleotide G at position 2 is deleted. The high deletion probability for position 4 means there is a 1% chance of a T insertion at this position (Table 2).

**Table 2:** Sequence-level probabilistic sequence defined by Equation (7)

| sequence | probability |
| --- | --- |
| $\mathcal{B}_1 = \text{CGAAT}$ | $a(1) = 0.9799$ |
| $\mathcal{B}_2 = \text{CAAT}$ | $a(2) = 0.01$ |
| $\mathcal{B}_3 = \text{CGATAT}$ | $a(3) = 0.01$ |
| $\mathcal{B}_4 = \text{CATAT}$ | $a(4) = 0.0001$ |

### 2.2 Constructing the probability sequence

#### 2.2.1 SAM files

In most next-generation sequencing applications, the estimated probability of sequencing error is quantified with the quality (or “Phred”) score attributed to each base call produced by sequencing instrument. The quality score *Q* is directly related to this estimated error probability: *ϵ* = 10^−*Q*/10^ (Ewing and Green, 1998), where *Q* typically ranges between 1 and 60 (with 60 being the lowest probability of error), depending on the sequencing platform and version of base-calling software. It is important to note that this quality score only measures the probability of error from the machine; 1 – *ϵ* is an estimate of the probability of no sequencing errors and does not account for any other source of error.

More formally, the probability that the base call is correct is expressed as:

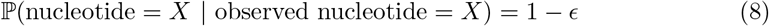

Unfortunately, quality scores have no information on the probabilities of the three other possible nucleotides if the base call is incorrect. In the absence of information about the other bases, we assume that these other probabilities are uniformly distributed.

Raw short read data are typically recorded in a FASTQ format that stores both the sequences (base calls) and base-specific quality scores. Since the reads often correspond to different positions of the target nucleic acid, *e.g*., randomly sheared genomic DNA, it is necessary to align the reads to identify base calls on different reads that represent the same genome position. This alignment step can be accomplished by mapping reads to a reference genome, by the *de novo* assembly of reads, or a hybrid approach that incorporates both methods. The aligned outputs are frequently recorded in the tabular Sequence Alignment/Map (SAM) format (Li et al., 2009a). Each row represents a short read, including the raw nucleotide sequence and quality strings; the optimal placement of the read with respect to the reference sequence (as an integer offset); and the compact idiosyncratic gapped alignment report (CIGAR) string, an application-specific serialization of the edit operations required to align the read to the reference. The SAM format contains much more information (https://samtools.github.io/hts-specs/SAMv1.pdf), but for our purposes we only need the placement, sequence, quality, and CIGAR string.

We employed the following procedure to construct the nucleotide-level probabilistic sequence from the contents of a SAM file. We initialize aligned sequence and quality strings with in all positions before the first read and after the last read, and ‘!’, which corresponds to a quality score of 0 (*Q* = 0), to all other positions. Next, we tokenize the CIGAR string into length-operation tuples, which determine how bases and quality scores from the raw strings are appended to the aligned versions. Deleted bases (‘D’ operations) are not assigned Phred scores, so we assume them to have 0 error probability.

Insertions (‘I’ operations) are non-trivial to include in the probabilistic sequence. Consider a short read with two bases inserted at position *j* (say, an `A` at position *j* + 1 and a `T` at position *j* + 2) and a short read with one insertion at position *j* (say, a `C`). It is entirely ambiguous whether the single insertion (`C`) aligns with the first insertion (`A`) or the second insertion (`T`) of the first short read. This is problematic for building up the matrix from reads aligned to the reference sequence. It is conceptually and computationally simpler to start from a populated matrix and sampling insertions. For our purposes, we only consider the alignment of these sequences with a reference sequence and thus do not consider insertions.

Some NGS platforms (*e.g*., Illumina) use paired-end reads where the same nucleic acid template is read in both directions. In these situations, we simply adjust all values by a factor of one half. For bases where the paired-end reads overlap, this has the effect of averaging the base probability 1 – *ϵ*. For example, if 1 – *ϵ* is 90% for `A` in one read and 95% `A` in its mate, then 0.925 is added to the `A` row in 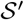 (with the remaining 0.075 uniformly distributed across the other nucleotides). If the two reads were 60% `A` and 55% `C` at the same position, then we would increment the corresponding column vector (`A, T, C, G`) by (0.6/2,0.1/2,0.1/2,0.1/2) for the first read and (0.15/2, 0.15/2, 0.55/2,0.15/2) for the second, resulting in an addition of (0.375, 0.125, 0.325, 0.125) for this pair. Bases outside of the overlapping region contribute a maximum of 0.5 to 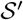, because the base call on the other read is missing data. This approach has the advantage of making the parsing of SAM files trivially parallelizable since we do not need to know how reads are paired. In addition, the coverage calculated from 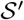 is scaled to the number of templates rather than the number of reads.

#### 2.2.2 Consensus sequence FASTQ files

Full length or partial genome sequences are now frequently the product of next-generation sequencing, by taking the consensus of the aligned or assembled read data. However, the original read data are often not published alongside the consensus sequence. Some consensus sequences are released in a format where the bases are annotated with quality scores, *e.g*., FASTQ. There are several programs that provide methods to convert a SAM file into a consensus FASTQ file (Li et al., 2004; Keith et al., 2002; Li et al., 2008). These programs use slightly different methods for generating consensus quality scores, but filter quality scores for the majority base. For example, suppose there are three reads with the following base calls at position *j*: `A` with *Q* = 30, `A` with *Q* = 31, and `C` with *Q* = 15. Calculation of the consensus quality score will thereby exclude the *Q* = 15 value and report a quality score calculated from *Q* = 30 and *Q* = 31, with the details of the calculation differing by software.

This omission makes it challenging for us to generate an 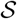 matrix from a consensus FASTQ file. Given the consensus base and its associated quality score at position *j*, we must assume that the other bases are all equally likely with probability *ϵ_j_*/3 (similar to Kuo et al. (2018) and Chapter 5 of Kozlov (2018)). For example, let’s assume the output sequence after fragment sequencing and alignment is `ACATG` and its associated quality scores are respectively *Q* = (60, 30, 50,10, 40). The probabilistic sequence is:

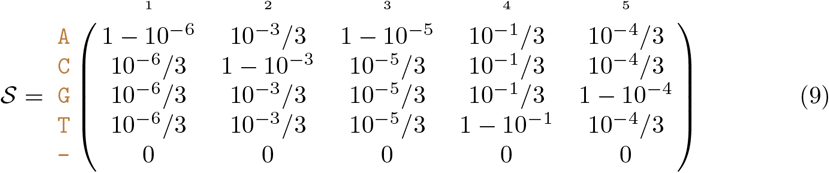

Usually, the genetic sequence `ACATG` would be considered as certain and quality scores discarded. In contrast, the probability of the sequence `ACATG` is only 0.899 within the probabilistic sequence framework.

Incorporating deletions in the absence of raw data is also challenging. If one is willing to assume a global deletion rate, then it is possible to extend the parameterization of 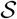. For example, if the probability of a single nucleotide deletion is *d*, then the probability of the called base is (1 – *d_j_*)(1 – *ϵ_j_*) and the other three nucleotides have probability (1 – *d*)*ϵ_j_*/3. Hence, if we assume the base call is `A`, the column of the nucleotide-level probabilistic sequence for that position is

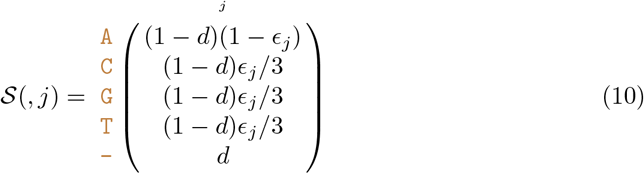

Since the FASTQ file only has a single sequence, we do have the same issues with alignment of differing lengths of insertions. In fact, insertions are only insertions relative to the reference sequence; they can simply be treated as observed nucleotides with an associated quality score. It would be possible to give insertions special treatment, however, by defining a global insertion rate. This insertion rate can be expressed as a deletion rate relative to the observed sequence, and thus one minus the insertion rate can be treated as the deletion rate in the probabilistic sequence. As with the deletion rate, this requires an assumption about a global rate which may be arbitrary.

#### 2.2.3 Consensus sequence FASTA files

If we do not have access to any base quality information, *e.g*., the consensus sequence is published as a FASTA file, then our ability to populate 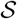 is severely limited. Any uncertainty that we impose upon the data will be a principled assumption. The error probability at the *j* position of the consensus sequence can be simulated as a beta distribution, i.e.,

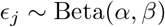

The called base at position *j* has probability 1 – *ϵ_j_*, and the remaining bases are assigned *ϵ_j_*/3. To incorporate deletions, another probability d can be generated as the *gap probability*. With these defined, the nucleotide-level probabilistic sequence at the *j*th column (assuming the base call at position *j* was A) can be written as above. This probabilistic sequence is completely fabricated, *i.e*., not based on any empirical data. However, the sensitivity of an analysis can be evaluated by choosing different values of *α*, *β*, and *d* (*e.g*., based on previous studies) and propagating these uncertainties into downstream analyses. The results from such an analysis would not indicate anything about the sequence itself but could be used to determine how robust the methods are to increased sequence uncertainty.

### 2.3 Propagation of uncertainty via resampling

The most general way to propagate uncertainty is through resampling. Given 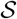 and assuming that individual nucleotides are independent outcomes — or pre-computing a reduced set of *m* sequences and calculating the distribution *a* of their joint probabilities, *i.e*., the sequence-level approach — we can propagate uncertainty by running downstream analyses on each set of sampled sequences.

At a nucleotide level, we are sampling from a multinomial distribution. If the *j*th column of 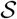 is (0.5, 0.2, 0.2, 0.09, 0.01), then we could sample `A` with 50% probability, `C` with 20%, etc. As with other sequence analyses, we can censor the positions that do not have enough coverage. We arbitrarily chose to censor any position that had fewer than 10 reads, which can be determined by summing the column of the probabilistic sequence.

In a maximum likelihood framework, this procedure is similar to bootstrapping. In fact, the ultimate effect of this is to decrease the bootstrap confidence to a level that is more in line with the measured uncertainty in the base calls.

In a Bayesian framework, the multinomial sampling could be incorporated as prior distribution on each nucleotide. For large collections of large sequences in an Markov Chain Monte Carlo algorithm, this increases the dimensionality dramatically.

### 2.4 Implementation

A C program has been written to convert SAM files into our matrix representation. The program assumes that the reads are aligned to a reference, then uses that reference to initiate the matrix. Because of our methods for handling paired reads, the program is able to stream the file line-by-line in a parallel computing environment.

The resampling algorithm defined above has been implemented in the R programming language. A shell script is used to repeatedly call the necessary R functions and apply the resampling algorithm to all outputs of the C program until the desired number of samples is obtained. All of the code for this project is available at https://github.com/Poonlab/SUP.

## 3 Applications

### 3.1 SARS-CoV-2 lineage assignment

In this section, we apply the re-sampling method to evaluate the impact of sequencing error on the lineage assignments of SARS-CoV-2. Sequences are sampled from 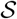 and then assigned a lineage based on the current state-of-the-art in phylogenetic analysis.

We use the lineage designation algorithm described in Rambaut et al. (2020) and assign our sequences to lineages using the pangoLEARN tool (Pangolin version 2.3.2, pangoLEARN version 2021-02-21) that the authors have made available (github.com/cov-lineages/Pangolin). This tool uses a decision tree model to determine which lineage a given sequence is most likely to belong to. We demonstrate that even the best available tools are underestimating the variance and therefore producing overconfident conclusions.

#### 3.1.1 Data

The data for this application were downloaded from NCBI’s SRA web interface (https://www.ncbi.nlm.nih.gov/sra/?term=txid2697049) on July 17th, 2021. Search results were filtered to only include records that had SAM files so that our alignments were consistent with the originating work. To select which runs to download, an arbitrary selection of 5-10 records from each of 20 non-sequential results pages were chosen. Once collecting the run accession numbers from the search results, an R script was run to download the relevant files and check that all information was complete. 23 out of 275 files were incomplete due to technical errors during the download process and a further 4 were rejected due to lack of CIGAR strings. The SRA accession numbers for the sequences we used are provided in the Appendix.

#### 3.1.2 Re-sampling the probabilistic sequence

Since pangoLEARN is a pre-trained model, assigning lineage designations to a large number of resampled genome sequences is not computationally burdensome. Sampling 5,000 different sequences from a probabilistic sequence can be done in a reasonable amount of time, even on a mid-range consumer laptop.

For this analysis we use the most basic resampling strategy described above. We sample base calls from the multinomial distribution then use pangoLEARN to determine the lineage assignment.

Figure 1 shows that the consensus sequence is almost always assigned to the same lineage as the majority of the resamples; the full results are in the Appendix. The proportion of resamples with the same lineage as the consensus sequence is very rarely 100% and can be as low as 32.86% (accession number ERR4440425). There were 52 cases where the proportion agreeing with the consensus sequence was either exactly 0 or less than 1%, and these cases occurred when the most common lineage sampled was labelled or “None” (sequences are labelled “None” when pangolin’s classification does not reach a confidence threshold). It is noteworthy that the only times where 100% of resampled sequences agreed are when the lineage call was “None” (13 cases) or for the lineage labelled B.1.1.7 (16 cases). This lineage represents 6% of our data and is a significantly more infectious lineage that is of special concern to health authorities (Wise, 2020; European Centre for Disease Prevention and Control, 2021).

**Figure 1:**
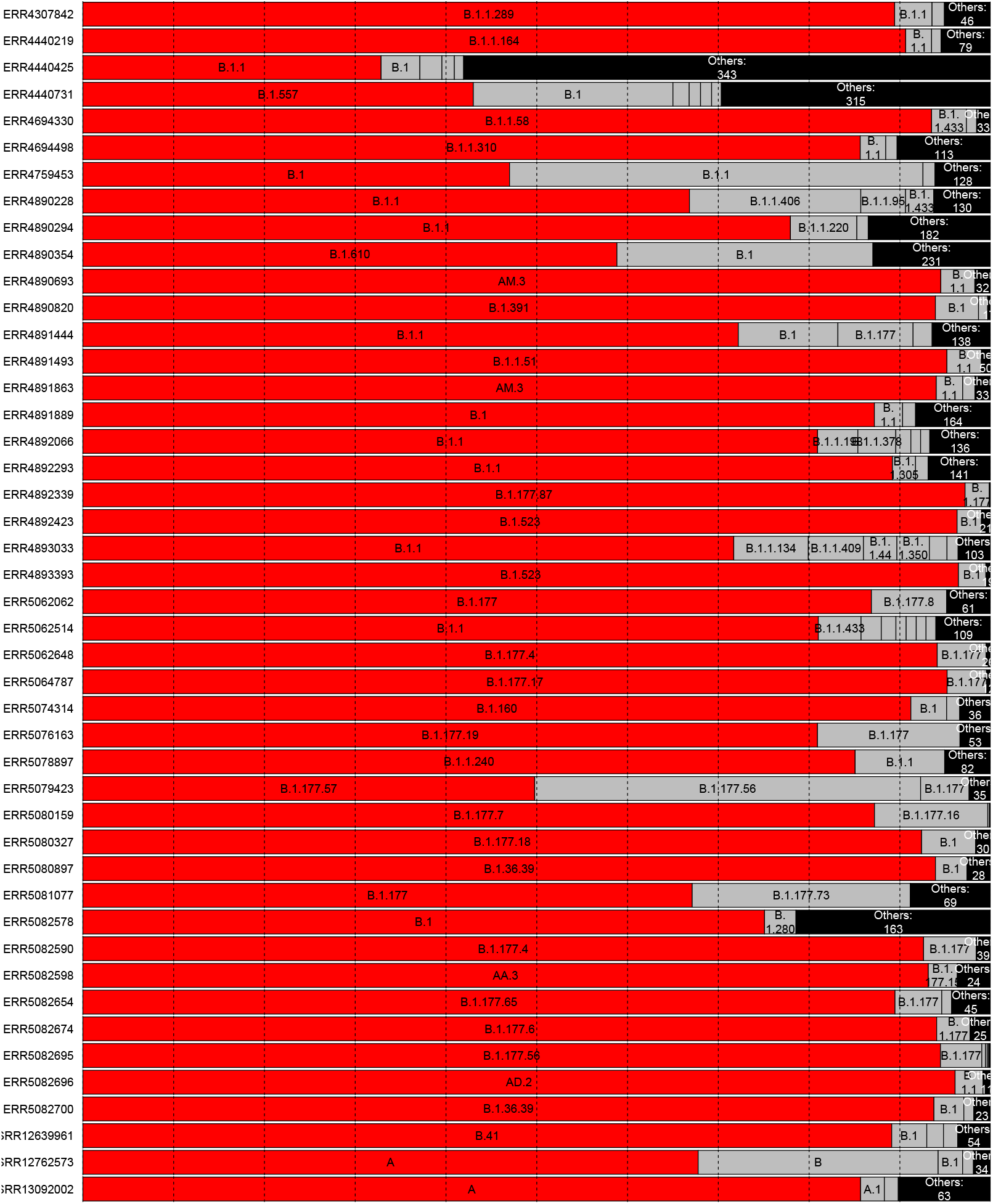
Visualization of called lineages from Pangolin. Red bars indicate the lineage of the most probable sequence and grey bars represent other sequences called from the same SAM file. Any lineage with fewer than 100 observations in the simulated sequences was grouped into the “Other” category. There were 95 sequences total, but we only the plotted ones where the second most common lineage designation had more than 250 observations.

**Figure 2:**
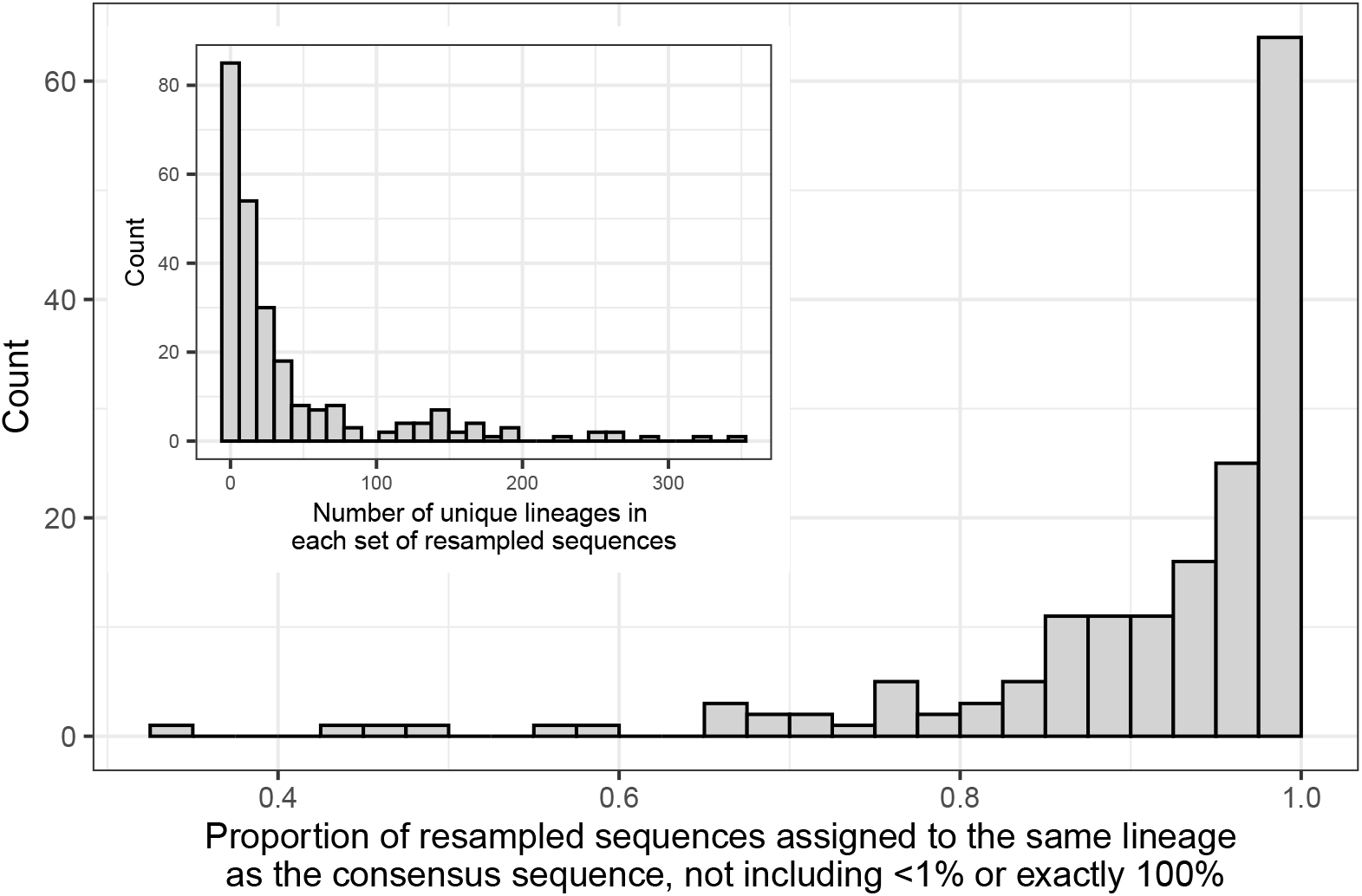
**Main plot:** Proportion of resampled sequences that are assigned to the same lineage as the consensus sequence. One proportion is calculated for each SAM file. The sets of resampled sequences where the proportion was less than 1% or exactly 100% are explained in Section 3.1.2. **Inset:** The number of distinct lineage assignments within each set of resampled sequences.

### 3.2 Clock rate estimation for SARS-CoV-2

The molecular clock rate (the number of mutations per site per unit of time) of a phylogenetic tree is found by considering both the number of mutations for each observed sequence relative to the root of the tree and the sample dates of those sequences. Assuming heterochronous sampling dates, the rate of mutations can be estimated by regressing the number of mutations against the sampling date. In the simplest case the clock rate is the slope estimate from a linear regression, which assumes a fixed clock rate. Polynomial and non-linear clock rates can be estimated (Sagulenko et al., 2018), as well as Bayesian non-parametric estimates (Drummond and Bouckaert, 2015).

The clock rate (slope of a root-to-tip regression) for SARS-CoV-2 is commonly estimated as a fixed rate near 0.001 mutations per site per year (Duchene et al., 2020; Choudhary et al., 2021; Song et al., 2021; Nie et al., 2020; Geidelberg et al., 2021). Using the same resampling methods as above, we estimate a clock rate for trees estimated from each of 50 resamples and for the tree estimated based on the reported consensus sequences.

To obtain the data, we sampled genomes uniformly from each month of recorded data in GenBank, using filters to ensure that the genomes were complete and had an associated SAM file. We further had to filter out SAM files that were incomplete or did not contain the CIGAR strings necessary for alignment, leaving us with 244 sequences. The associated SRA accession numbers are provided in the Appendix.

Our re-sampling method will, by definition, introduce other possible mutations beyond what the consensus sequence suggests. Because of this, the apparent number of mutations between a re-sampled genome and the estimated root is a function of the coverage, with more positions read or more uncertainty in the sequence leading to artificially inflated terminal branch lengths. Furthermore, we are sampling nucleotides at each position independently of other positions as well as independently of ancestral sequences. This implies that the estimates of the time for the most recent common ancestor are not reliable. However, assuming that the sequences have comparable levels of uncertainty, each branch increases by a similar amount and the clock rate should not be affected, only the intercept.

The sequences that we acquired did not have comparable levels of uncertainty; the viruses sampled early in the pandemic had considerably higher uncertainty, most likely due to a lack of consistent laboratory guidelines for sequencing this new virus. To account for this, we calculated the sum of 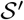 for each sequence and applied Statistical Process Control techniques to ensure that all of the sequences had a similar level of coverage. In particular, we calculated the mean coverage of the sequences in our data set, 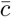, and the standard deviation of the coverage, *s*. We removed any sequences outside of 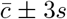, recalculated 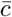 and *s*, and iterated the removal process until all sequence coverages were within the bounds, amounting to 20 removed sequences.

The clock rate was estimated using TreeTime (Sagulenko et al., 2018). We recorded the clock rate and standard error from the time tree constructed using the consensus sequences and compared this to the clock rate and standard deviations of the estimated clock rates in the resampled sequences. The tree built from consensus sequences had a clock rate of 6.5 × 10^−4^ with a standard error of 8.01 × 10^-5^. The mean of the clock rates for all of the sets of resampled sequences was 8.6 × 10^-4^ with standard deviation of 5.3 × 10^-4^, which is approximately 1.6 times as large as the standard error for the consensus sequences.

The estimates of the clock rate are shown in Figure 3. The red line and shaded region are the clock rate for the tree built from consensus sequences along with ±1.96 standard errors. Rate estimates from Duchene et al. (2020) (n=122), Choudhary et al. (2021) (n=261), Song et al. (2021) (n=29), Nie et al. (2020) (n=112), and Geidelberg et al. (2021) (n=77) are also labelled on the plot with purple error bars for 95% Bayesian Credible Intervals (BCI) or 95% Highest Posterior Density (HPD), indicating that the rates and errors from each root-to-tip regression are in line with other published results. Figure 3 demonstrates that the estimated evolutionary rates have an average close to the rate estimated from our tree estimated from consensus sequences as well as the rates from other studies, but each of the individual error bars (from the five studies identified above) miss the excess variation due to sequence uncertainty.

**Figure 3:**
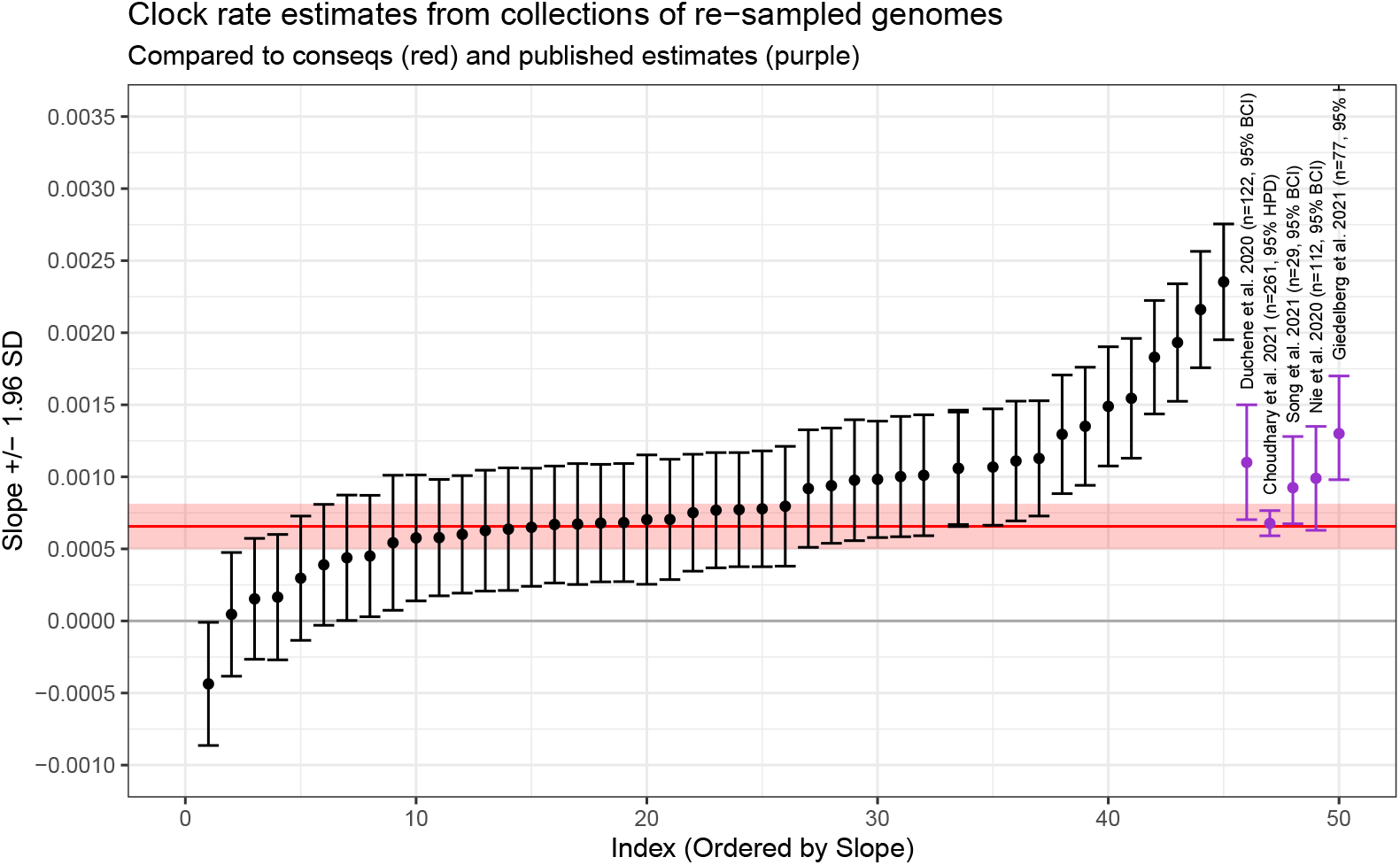
Clock rates (slope) and 95% Confidence Intervals for the collections of re-sampled sequences. The red line and red shaded region are the clock rate and 95% CI for the consensus sequences. The purple points and error bars are the clock rates and error intervals (either Bayesian Credible Interval or Highest Posterior Probability) from published studies, as labelled. The re-sampled sequences are in line with the consensus sequences as well as the published sequences, but represent a much larger variation due to the uncertainty in the original genome sequences.

## 4 Conclusions

The files produced by NGS platforms include valuable information about the quality of base calls which should be propagated into analyses. In this study, we have demonstrated that these errors in base calling can lead to different conclusions when determining a lineage via Pangolin and that the variance in clock rate estimates is larger than previously shown due to these errors. Both of these situations could lead to incorrect conclusions, such as missing a variant of interest or making overconfident conclusions about the date of the first case of COVID-19. The potential for errors in base calls should always be taken into account when making decisions based on genetic sequencing data.

Our analysis of Pangolin lineage classification demonstrates that the uncertainty in the base calls has a non-trivial effect on the potential lineage calls. The reported lineage classifications are based on a sophisticated classification algorithm which has high confidence in the predicted category, but this assumes that the input sequence is known without error. We are not aware of any classification system that incorporates per-base error, so we suggest that interpretations of the output of any classification system be interpreted with reference to the uncertainty in their sequence.

Our clock rate estimation suggest that the confidence/credible intervals for the published clock rates are underestimated. As with lineage classification, we are not aware of any clock rate estimation procedures that incorporate the uncertainty in the base calls of the sequences. Researchers should be conscious of this potential source of currently unacknowledged error when reporting any results from sequenced genomes.

## 5 Discussion

The primary contribution of this research is the construction of the probability sequence, which allows for a wide variety of future research directions. The direction we described here is focused on re-sampling, which allows a more complete appraisal of the variance in the estimates (or provides a reasonable prior distribution in a Bayesian setting), while comparing results for the most likely sequences provide a measure of robustness to sequence uncertainty.

In the absence of SAM files, FASTQ files for the consensus sequence also offer quantification of the potential sequence read errors. We have already shown that probability sequences can be trivially constructed by spreading the remaining error probability uniformly across the remaining bases. Another use of FASTQ files would be to construct a probability sequence as a reference genome for a given category. This would entail collecting all available FASTQ files for a given lineage designation and using them in the construction of a probability sequence as if they were short reads in a SAM file. From here, lineage designation for a newly acquired sequence (and its probability sequence) could be performed via a hypothesis test for whether the probability sequences are sufficiently similar.

Our proposed methods can result in a linear increase in computational expense. Even the method based on ordering the sequences by likelihood inevitably requires re-running the analysis numerous times. However, we have demonstrated that the uncertainty in the sequences themselves can lead to major changes to the interpretations of the results. The so-called “consensus sequence” is simply the most likely sequence, and the reported uncertainty is not merely an academic curiosity. Ideally individual analyses would be constructed to take nucleotide-level uncertainty into account. For instance, phylogenies have been estimated based on uncertain sequence information in Ross and Markowetz (2016); Jahn et al. (2016); Zafar et al. (2017) but the uncertainty is not derived from base quality scores. An extension of these methods to incorporate the base quality scores may be a worthwhile research direction.

Computational burden can also be reduced by sorting the sequences in decreasing uncertainty. By doing so, we can analyse how the increases in uncertainty affect our analysis. It is possible to devise an algorithm that puts the sequences in order of their uncertainty without calculating the uncertainty for every sequence. The consensus sequence is clearly the least uncertain sequence, and by the construction of the probability sequence, it is computationally easy to calculate sequence uncertainty relative to this sequence. Because of this, it should be reasonable to calculate all sequence uncertainties for a single mutation, then all pairs of mutations, and so on (we note that it is possible for some pairs of mutations to be more likely than single mutations, thus this algorithm cannot readily calculate the *n* most likely sequences). This allows for easy sorting of the most likely sequences, assuming enough sequence uncertainties have been calculated. With this sorted list, we can examine each sequence in increasing uncertainty in order to approximate the distribution of sequence uncertainties. Any model that uses sequence data could be re-fit with each sequence to investigate the robustness of that model to sequence uncertainty.

Our analysis focused on lineage classification according to the Pangolin model as well as estimation of the clock rate. The importance of incorporating sequence uncertainty is not confined to these applications; any analysis involving sequenced genomes would benefit from some method of incorporating the uncertainty or including some measure of robustness. For example, the estimated frequency of alleles in the population could be used as the probability sequence, then propagated into further analysis.

Our method does not preclude tertiary analyses to test for systematic errors. For instance, De Maio et al. (2020) suggest that some errors arise due to issues in the sequencing protocol in particular laboratories. Our method allows for adjustments of the base call quality score, such as in Brockman et al. (2008), correcting for laboratory-specific errors, as well as more sophisticated definitions of genome likelihoods (*e.g*., Li et al., 2004; DePristo et al., 2011; Li et al., 2009b).

We have evaluated an algorithm to include insertion events in a re-sampling scheme, but many of the resultant sequences were not mappable to known sequences. The Pangolin lineage assignment system appears to treat insertions differently from single nucleotide polymorphisms, and our method of sampling insertions is incompatible with their treatment of insertions in lineage assignment operations. This is potentially because the sampled base pair at any given position is independent of each other position, and the insertions observed in real-world data are possibly always associated with particular mutations elsewhere. However, insertions in the SARS-CoV-2 genome have been relatively rare.

This study should not be taken in any way as a criticism of the Pangolin lineage assignment procedure. Rather, Pangolin was chosen as it is the state-of-the art tool for lineage classification. The phylogeny created by this team has been a vital resource for researchers and for public health professionals. In particular, the PANGO label for the current Variants of Concern (VOCs), especially B.1.1.7, are the labels being used worldwide by news organizations. The output from Pangolin and many other bioinformatics tools are usually interpreted as *deterministic* results. This study is an argument that inherent uncertainty in sequencing warrants propagation into downstream analyses.

**Table 3:** Accession numbers for the resampling application. The prefix ERR indicates that the sequence comes from the European Nucleotide Archive, whereas the prefix SRR indicates that it comes from the NCBI’s Short Read Archive.

|  |  |  |  |  |  |
| --- | --- | --- | --- | --- | --- |
| ERR4085809 | ERR4890354 | ERR4892048 | ERR5064166 | ERR5082590 | SRR13021020 |
| ERR4204823 | ERR4890371 | ERR4892066 | ERR5064294 | ERR5082598 | SRR13021022 |
| ERR4305816 | ERR4890386 | ERR4892112 | ERR5064346 | ERR5082599 | SRR13021027 |
| ERR4307842 | ERR4890403 | ERR4892152 | ERR5064787 | ERR5082600 | SRR13021032 |
| ERR4440194 | ERR4890427 | ERR4892200 | ERR5064811 | ERR5082606 | SRR13021033 |
| ERR4440219 | ERR4890531 | ERR4892203 | ERR5074314 | ERR5082610 | SRR13021035 |
| ERR4440247 | ERR4890572 | ERR4892293 | ERR5076163 | ERR5082622 | SRR13021038 |
| ERR4440332 | ERR4890609 | ERR4892339 | ERR5076748 | ERR5082630 | SRR13021042 |
| ERR4440354 | ERR4890693 | ERR4892386 | ERR5077151 | ERR5082645 | SRR13021047 |
| ERR4440373 | ERR4890746 | ERR4892392 | ERR5077411 | ERR5082654 | SRR13021052 |
| ERR4440402 | ERR4890771 | ERR4892423 | ERR5077618 | ERR5082656 | SRR13021053 |
| ERR4440425 | ERR4890819 | ERR4893013 | ERR5077713 | ERR5082673 | SRR13021059 |
| ERR4440731 | ERR4890820 | ERR4893031 | ERR5077924 | ERR5082674 | SRR13021061 |
| ERR4692420 | ERR4890881 | ERR4893033 | ERR5078210 | ERR5082694 | SRR13021067 |
| ERR4692568 | ERR4890926 | ERR4893037 | ERR5078863 | ERR5082695 | SRR13021072 |
| ERR4692877 | ERR4890974 | ERR4893080 | ERR5078897 | ERR5082696 | SRR13021073 |
| ERR4692945 | ERR4891001 | ERR4893138 | ERR5079000 | ERR5082700 | SRR13021077 |
| ERR4693014 | ERR4891011 | ERR4893184 | ERR5079423 | ERR5082702 | SRR13021084 |
| ERR4693019 | ERR4891037 | ERR4893186 | ERR5079699 | ERR5082706 | SRR13021090 |
| ERR4693495 | ERR4891061 | ERR4893197 | ERR5080131 | ERR5082708 | SRR13021093 |
| ERR4693801 | ERR4891103 | ERR4893242 | ERR5080159 | ERR5082710 | SRR13021097 |
| ERR4693865 | ERR4891178 | ERR4893353 | ERR5080327 | ERR5082711 | SRR13021098 |
| ERR4694010 | ERR4891235 | ERR4893393 | ERR5080504 | ERR5082712 | SRR13021099 |
| ERR4694330 | ERR4891238 | ERR4999251 | ERR5080893 | SRR11433882 | SRR13021104 |
| ERR4694380 | ERR4891261 | ERR4999255 | ERR5080897 | SRR11433888 | SRR13021107 |
| ERR4694400 | ERR4891304 | ERR4999275 | ERR5080913 | SRR11433893 | SRR13021109 |
| ERR4694498 | ERR4891415 | ERR4999282 | ERR5080918 | SRR12639958 | SRR13021111 |
| ERR4694556 | ERR4891433 | ERR5060778 | ERR5081077 | SRR12639961 | SRR13021113 |
| ERR4694571 | ERR4891444 | ERR5062004 | ERR5081293 | SRR12749715 | SRR13021115 |
| ERR4694617 | ERR4891493 | ERR5062062 | ERR5081301 | SRR12749716 | SRR13021124 |
| ERR4759453 | ERR4891497 | ERR5062388 | ERR5081304 | SRR12762573 | SRR13021130 |
| ERR4869446 | ERR4891532 | ERR5062514 | ERR5081316 | SRR13020989 | SRR13021131 |
| ERR4869458 | ERR4891572 | ERR5062571 | ERR5081322 | SRR13020990 | SRR13021133 |
| ERR4869480 | ERR4891675 | ERR5062648 | ERR5081836 | SRR13020991 | SRR13021134 |
| ERR4869487 | ERR4891711 | ERR5062729 | ERR5082214 | SRR13020998 | SRR13021135 |
| ERR4869497 | ERR4891715 | ERR5062935 | ERR5082346 | SRR13020999 | SRR13021143 |
| ERR4890228 | ERR4891805 | ERR5063143 | ERR5082556 | SRR13021003 | SRR13092002 |
| ERR4890271 | ERR4891841 | ERR5063165 | ERR5082561 | SRR13021008 | SRR13592146 |
| ERR4890285 | ERR4891863 | ERR5063539 | ERR5082569 | SRR13021010 |  |
| ERR4890294 | ERR4891889 | ERR5063807 | ERR5082576 | SRR13021011 |  |
| ERR4890337 | ERR4891898 | ERR5063813 | ERR5082578 | SRR13021013 |  |
| ERR4890352 | ERR4891988 | ERR5063922 | ERR5082580 | SRR13021017 |  |

**Table 4:** Accession numbers used in the root-to-tip application. The prefix ERR indicates that the sequence comes from the European Nucleotide Archive, whereas the prefix SRR indicates that it comes from the NCBI’s Short Read Archive.

|  |  |  |  |  |  |
| --- | --- | --- | --- | --- | --- |
| ERR4333012 | ERR4598849 | ERR4645575 | ERR4763502 | ERR4893972 | ERR5189526 |
| ERR4422411 | ERR4599609 | ERR4647168 | ERR4763917 | ERR4905630 | ERR5196271 |
| ERR4423341 | ERR4632053 | ERR4651160 | ERR4788047 | ERR4906265 | ERR5240413 |
| ERR4423907 | ERR4633143 | ERR4652052 | ERR4792808 | ERR4989718 | ERR5277731 |
| ERR4424340 | ERR4633439 | ERR4652877 | ERR4793403 | ERR5020396 | ERR5293387 |
| ERR4424995 | ERR4633631 | ERR4659368 | ERR4824581 | ERR5021500 | ERR5304264 |
| ERR4437121 | ERR4637060 | ERR4667806 | ERR4824816 | ERR5025790 | ERR5307708 |
| ERR4437452 | ERR4639632 | ERR4668406 | ERR4824949 | ERR5027044 | ERR5314268 |
| ERR4438147 | ERR4639682 | ERR4668440 | ERR4825023 | ERR5040813 | ERR5316846 |
| ERR4438486 | ERR4639875 | ERR4668990 | ERR4826433 | ERR5041080 | ERR5334211 |
| ERR4459703 | ERR4640333 | ERR4669218 | ERR4827255 | ERR5052912 | ERR5339018 |
| ERR4459988 | ERR4640467 | ERR4686528 | ERR4835054 | ERR5052951 | ERR5339175 |
| ERR4460366 | ERR4640673 | ERR4686632 | ERR4848811 | ERR5054027 | ERR5349715 |
| ERR4463279 | ERR4641513 | ERR4686760 | ERR4849464 | ERR5057371 | ERR5353067 |
| ERR4581201 | ERR4641559 | ERR4688435 | ERR4860936 | ERR5058153 | ERR5379217 |
| ERR4584371 | ERR4642761 | ERR4688535 | ERR4861339 | ERR5177079 | SRR12349113 |
| ERR4584814 | ERR4643065 | ERR4699831 | ERR4874605 | ERR5181042 | SRR12349131 |
| ERR4597698 | ERR4643184 | ERR4706873 | ERR4874858 | ERR5187190 |  |
| ERR4597906 | ERR4644945 | ERR4763252 | ERR4878261 | ERR5188799 |  |

## Appendix

The following table lists all of the NCBI SRA accession numbers used in this paper.

## References

Beerenwinkel, N. and Zagordi, O. (2011). Ultra-deep sequencing for the analysis of viral populations. Current Opinion in Virology, 1(5):413–418.

Brockman, W., Alvarez, P., Young, S., Garber, M., Giannoukos, G., Lee, W. L., Russ, C., Lander, E. S., Nusbaum, C., and Jaffe, D. B. (2008). Quality scores and SNP detection in sequencing-by-synthesis systems. Genome Research, 18(5):763–770.

Choudhary, M. C., Crain, C. R., Qiu, X., Hanage, W., and Li, J. Z. (2021). Severe Acute Respiratory Syndrome Coronavirus 2 (SARS-CoV-2) Sequence Characteristics of Coronavirus Disease 2019 (COVID-19) Persistence and Reinfection. Clinical Infectious Diseases, (ciab380).

Clement, N. L., Snell, Q., Clement, M. J., Hollenhorst, P. C., Purwar, J., Graves, B. J., Cairns, B. R., and Johnson, W. E. (2010). The GNUMAP algorithm: Unbiased probabilistic mapping of oligonucleotides from next-generation sequencing. Bioinformatics, 26(1):38–45.

De Maio, N., Walker, C., Borges, R., Weilguny, L., Slodkowicz, G., and Goldman, N. (2020). Issues with SARS-CoV-2 sequencing data. https://virological.org/t/issues-with-sars-cov-2-sequencing-data/473, Accessed 2021-11-24.

DePristo, M. A., Banks, E., Poplin, R., Garimella, K. V., Maguire, J. R., Hartl, C., Philippakis, A. A., del Angel, G., Rivas, M. A., Hanna, M., McKenna, A., Fennell, T. J., Kernytsky, A. M., Sivachenko, A. Y., Cibulskis, K., Gabriel, S. B., Altshuler, D., and Daly, M. J. (2011). A framework for variation discovery and genotyping using next-generation DNA sequencing data. Nature Genetics, 43(5):491–498.

Doronina, N. V. (2005). Phylogenetic position and emended description of the genus Methylovorus. INTERNATIONAL JOURNAL OF SYSTEMATIC AND EVOLUTIONARY MICROBIOLOGY, 55(2):903–906.

Drummond, A. J. and Bouckaert, R. R. (2015). Bayesian Evolutionary Analysis with BEAST. Cambridge University Press.

Duchene, S., Featherstone, L., Haritopoulou-Sinanidou, M., Rambaut, A., Lemey, P., and Baele, G. (2020). Temporal signal and the phylodynamic threshold of SARS-CoV-2. Virus Evolution, 6(2).

European Centre for Disease Prevention and Control (2021). SARS-CoV-2 variants of concern as of 26 November 2021. https://www.ecdc.europa.eu/en/covid-19/variants-concern, Accessed 2021-11-26.

Ewing, B. and Green, P. (1998). Base-Calling of Automated Sequencer Traces Using *Phred*. II. Error Probabilities. Genome Research, 8(3):186–194.

Fuller, C. W., Middendorf, L. R., Benner, S. A., Church, G. M., Harris, T., Huang, X., Jovanovich, S. B., Nelson, J. R., Schloss, J. A., Schwartz, D. C., and Vezenov, D. V. (2009). The challenges of sequencing by synthesis. Nature Biotechnology, 27(11):1013–1023.

Fumagalli, M., Vieira, F. G., Korneliussen, T. S., Linderoth, T., Huerta-Sánchez, E., Albrechtsen, A., and Nielsen, R. (2013). Quantifying Population Genetic Differentiation from Next-Generation Sequencing Data. Genetics, 195(3):979–992.

Geidelberg, L., Boyd, O., Jorgensen, D., Siveroni, I., Nascimento, F. F., Johnson, R., Ragonnet-Cronin, M., Fu, H., Wang, H., Xi, X., Chen, W., Liu, D., Chen, Y., Tian, M., Tan, W., Zai, J., Sun, W., Li, J., Li, J., Volz, E. M., Li, X., and Nie, Q. (2021). Genomic epidemiology of a densely sampled COVID-19 outbreak in China. Virus Evolution, 7(1).

Gompert, Z. and Buerkle, C. A. (2011). A Hierarchical Bayesian Model for Next-Generation Population Genomics. Genetics, 187(3):903–917.

Goodwin, S., McPherson, J. D., and McCombie, W. R. (2016). Coming of age: Ten years of next-generation sequencing technologies. Nature Reviews Genetics, 17(6):333–351.

Jahn, K., Kuipers, J., and Beerenwinkel, N. (2016). Tree inference for single-cell data. Genome Biology, 17(1):86.

Keith, J. M., Adams, P., Bryant, D., Kroese, D. P., Mitchelson, K. R., Coachran, D. A. E., and Lala, G. H. (2002). A simulated annealing algorithm for finding consensus sequences. Bioinformatics, 18(11):1494–1499.

Kozlov, O. (2018). Models, Optimizations, and Tools for Large-Scale Phylogenetic Inference, Handling Sequence Uncertainty, and Taxonomic Validation.

Kuhner, M. K. and McGill, J. (2014). Correcting for Sequencing Error in Maximum Likelihood Phylogeny Inference. G3 Genes—Genomes—Genetics, 4(12):2545–2552.

Kuo, T., Frith, M. C., Sese, J., and Horton, P. (2018). EAGLE: Explicit Alternative Genome Likelihood Evaluator. BMC Medical Genomics, 11(2):28.

Li, H., Handsaker, B., Wysoker, A., Fennell, T., Ruan, J., Homer, N., Marth, G., Abecasis, G., and Durbin, R. (2009a). The Sequence Alignment/Map format and SAMtools. Bioinformatics, 25(16):2078–2079.

Li, H., Ruan, J., and Durbin, R. (2008). Mapping short DNA sequencing reads and calling variants using mapping quality scores. Genome Research, 18(11):1851–1858.

Li, M., Nordborg, M., and Li, L. M. (2004). Adjust quality scores from alignment and improve sequencing accuracy. Nucleic Acids Research, 32(17):5183–5191.

Li, R., Li, Y., Fang, X., Yang, H., Wang, J., Kristiansen, K., and Wang, J. (2009b). SNP detection for massively parallel whole-genome resequencing. Genome Research, 19(6):1124–1132.

NC-IUB (1986). Nomenclature for incompletely specified bases in nucleic acid sequences. Recommendations 1984.Nomenclature Committee of the International Union of Biochemistry (NC-IUB). Proceedings of the National Academy of Sciences of the United States of America, 83(1):4–8.

Nie, Q., Li, X., Chen, W., Liu, D., Chen, Y., Li, H., Li, D., Tian, M., Tan, W., and Zai, J. (2020). Phylogenetic and phylodynamic analyses of SARS-CoV-2. Virus Research, 287:198098.

O’Rawe, J. A., Ferson, S., and Lyon, G. J. (2015). Accounting for uncertainty in DNA sequencing data. Trends in Genetics, 31(2):61–66.

Rambaut, A., Holmes, E. C., O’Toole, A., Hill, V., McCrone, J. T., Ruis, C., du Plessis, L., and Pybus, O. G. (2020). A dynamic nomenclature proposal for SARS-CoV-2 lineages to assist genomic epidemiology. Nature Microbiology.

Richterich, P. (1998). Estimation of Errors in “Raw” DNA Sequences: A Validation Study. Genome Research, 8(3):251–259.

Robasky, K., Lewis, N. E., and Church, G. M. (2014). The role of replicates for error mitigation in next-generation sequencing. Nature Reviews Genetics, 15(1):56–62.

Ross, E. M. and Markowetz, F. (2016). OncoNEM: Inferring tumor evolution from single-cell sequencing data. Genome Biology, 17(1):69.

Sagulenko, P., Puller, V., and Neher, R. A. (2018). TreeTime: Maximum-likelihood phylodynamic analysis. Virus Evolution, 4(1):vex042.

Salk, J. J., Schmitt, M. W., and Loeb, L. A. (2018). Enhancing the accuracy of next-generation sequencing for detecting rare and subclonal mutations. Nature reviews. Genetics, 19(5):269–285.

Schneider, T. D. (2002). Consensus Sequence Zen. Applied bioinformatics, 1(3):111–119.

Schneider, T. D. and Stephens, R. (1990). Sequence logos: A new way to display consensus sequences. Nucleic Acids Research, 18(20):6097–6100.

Song, N., Cui, G.-L., and Zeng, Q.-L. (2021). Genomic Epidemiology of SARS-CoV-2 From Mainland China With Newly Obtained Genomes From Henan Province. Frontiers in Microbiology, 12:673855.

Stormo, G. D., Schneider, T. D., Gold, L., and Ehrenfeucht, A. (1982). Use of the ‘perceptron’ algorithm to distinguish translational initiation sites in E. coli. Nucleic Acids Research, 10(9):16.

Wise, J. (2020). Covid-19: New coronavirus variant is identified in uk. BMJ, 371.

Wu, S. H., Schwartz, R. S., Winter, D. J., Conrad, D. F., and Cartwright, R. A. (2017). Estimating error models for whole genome sequencing using mixtures of Dirichlet-multinomial distributions. Bioinformatics, 33(15):2322–2329.

Zafar, H., Tzen, A., Navin, N., Chen, K., and Nakhleh, L. (2017). SiFit: Inferring tumor trees from single-cell sequencing data under finite-sites models. Genome Biology, 18(1):178.

